# Protozoa populations are ecosystem engineers that shape prokaryotic community structure and function of the rumen microbial ecosystem

**DOI:** 10.1101/2020.05.15.080218

**Authors:** Ronnie Solomon, Tanita Wein, Bar Levy, Shahar Eshed, Rotem Dror, Veronica Reiss, Tamar Zehavi, Ori Furman, Itzhak Mizrahi, Elie Jami

**Author notes:** Corresponding author: Elie Jami, Agricultural Research Organization - the Volcani Center, 68 HaMaccabim Road, P.O.B 15159 Rishon LeZion 7505101, Israel.

## Abstract

Unicellular eukaryotes are an integral part of many microbial ecosystems communities where they are known to interact with their surrounding prokaryotic community – either as predators or as a mutualistic habitat. Within the rumen, one of the most complex host-associated microbial habitats, ciliate protozoa represent the main micro-eukaryotes, accounting for up to 50% of the microbial biomass. Nonetheless, the extent of the ecological effect of protozoa on the microbial community and on the rumen metabolic output remains largely understudied. To assess the role of protozoa on the rumen ecosystem, we established an *ex-vivo* system in which distinct protozoa sub-communities were introduced to native rumen prokaryotic community. We show that the different protozoa communities exert a strong and differential impact on the composition of the prokaryotic community, as well as its function including methane production. Furthermore, the presence of protozoa increases prokaryotic diversity with a differential effect on specific bacterial populations such as Gammaproteobacteria, *Prevotella* and Spirochetes. Our results suggest that protozoa mitigate the effect of competitive exclusion between bacterial species, thereby contributing to the maintenance of prokaryotic diversity in the rumen. Our findings put forward the rumen protozoa populations as potentially important ecosystem engineers for future microbiome modulation strategies.

## Introduction

Microbial community assemblages are determined by both abiotic and biotic factors driven by the environmental conditions and a complex network of microbial interactions between diverse microorganisms. These interactions include mutualism, competition, and predation that ultimately define the microbial community composition.

Microbial eukaryotes - such as protists - are ubiquitous in a wide range of environments and are known to play a pivotal role in regulating microbial community structure and function as well as their physicochemical environment (e.g., Pernthaler 2005 [1]). Protist-bacteria interactions range from mutualistic (e.g., metabolic exchange or scavenging of toxic compounds) [2–4], to antagonistic interplay that mainly comprises predation [1–3]. As predators, they are considered a major cause of bacterial mortality in microbial ecosystems, where they exert a top down control that was shown to greatly impact surrounding prey species. Protist-predation further leads to changes at the microbial community structure level, as well as at the single cell level, which can promote changes in bacterial morphology and evolution [1, 5–8].

In addition to their predatory interactions with their neighboring prokaryotes, protists were shown to positively interact with bacteria and archaea. This is exemplified by various interspecies exchanges of metabolites with the surrounding prokaryotic community [4, 9] as well as evidence of physical interaction with bacteria and archaea localized inside or surface attached prokarya contributing to their respective fitness [2, 10].

In host-associated microbial communities, protists were suggested to play a beneficial role for their host. For example, in the rhizosphere of plants, predation by protists promotes changes in microbial composition that accelerates nutrient cycling and supports the removal of pathogenic species [11–13]. Though less evidence exists regarding mammalian hosts, the protist community was suggested to confer protection to the host via immune mediated response against pathogens [14, 15]. Additionally, based on population analysis of the microbiome in the human gut, the presence of *Blastocystis* was suggested to play a critical role in maintaining bacterial diversity in the gut [15]. Such evidence suggests that protists not only play a crucial role for microbial community assemblages, but may also have a direct effect on the host species and alterations of these interactions have the potential to greatly affect microbial ecosystems. Nonetheless, the complex role of protists in diverse environments including animal-associated microbiomes remains largely understudied.

One of the most densely populated gut environments in which protozoa species are ubiquitously found is the upper digestive tract of ruminants termed rumen, where protozoa are estimated to encompass between 25% to 50% of the microbial biomass [16–18]. As an ecosystem, the rumen hosts one of the most complex microbial communities comprising bacteria, archaea and microbial eukaryotes, the latter being dominated by ciliate protozoa [19]. The protozoa population, like bacteria and archaea, encompass a large array of diverse species in the rumen environment [20], historically characterized by different morphologies and sizes, with species larger than 100 μm to smaller 10 μm [21]. Rumen protozoa are part of a complex microbial community responsible for the breakdown of plant feed into digestible molecules for the animal and accordingly, ruminant productivity has been, in recent years, tightly linked with the rumen microbial community composition [22–26]. Unlike bacteria, protozoa are not obligatory in the rumen and can be stably removed with no apparent ill-effect to the host, via defaunation [17].

Defaunations thus allows for controlled *in-vivo* experiments in which the role of the protozoa can be evaluated by either addition or subtraction of the whole ciliate protozoa community or specific subpopulations [17]. Such experiments revealed that the removal of protozoa from the rumen carries a tremendous effect on the production of metabolic end-products and nitrogen metabolism and that different protozoa taxa contribute differently to the metabolic aspects of the rumen parameters and animal physiology [17, 27–31]. Specifically, the absence of protozoa was shown to decrease methane emission in defaunated animals. This observation suggests a metabolic interaction between protozoa and methanogenic archaea, the sole producers of methane in the rumen. Protozoa are known for their production of hydrogen while the majority of methanogens in the rumen produce methane via the hydrogenotrophic pathway. This notion is reinforced by observations of physical association between protozoa and methanogens [4, 32–34].

Despite the importance of protozoa communities on the rumen ecosystem and our environment, the direct effect of protozoa community composition on the prokaryotic community structure and function was never examined in a controlled and defined experimental setup. To this end, we utilized an experimental setup in which different rumen protozoa sub-populations were established and exposed to an identical freshly sampled rumen prokaryotic community to characterize the resulting metabolic output as well as the microbial prokaryotic community dynamics. Our findings suggest that rumen protozoa play a central role in defining characteristics of the rumen ecosystem shaping rumen microbiome structure and metabolism.

## Materials and methods

### Animal Handling and Sampling

The experimental procedures used in this study were approved by the Faculty Animal Policy and Welfare Committee of the Agricultural Research Organization Volcani Research Center approval no. 737/17 IL, in accordance with the guidelines of the Israel Council for Animal Care.

Rumen fluid was sampled from three cows kept under the same diet for at least two month and transferred immediately to an oxygen free environment in an anaerobic glove box for further processing.

### Protozoa separation

In order to obtain different populations of protozoa, the rumen samples underwent a series of size filtration and washings similar to the procedure performed in [32]. Briefly, the rumen fluid was mixed in a 1:1 ratio with a warmed, anaerobic Coleman buffer (Williams and Coleman, 2012), and incubated in a separating funnel for 1 h under anaerobic conditions at 39°C. The settled protozoa fraction was transferred to a fresh tube with warm Coleman buffer. Prior to filtration a subset of the whole protozoa community was put aside and represents the all protozoa group in the study. The rest of the protozoa underwent consecutive filtration using nylon net filters (Merck Millipore, Darmstadt, Germany) of different sizes (i.e., 100 μm, 60 μm, 40 μm, 10 μm). The retentate on each filter and the filtrate of the last 10 μm filtering were then washed with a warm anaerobic Coleman buffer [20]. A subset of each fraction was taken for counting under light microscopy in order to be able to inoculate the microcosms with the same number of protozoa. Paraformaldehyde at a final volume of 4% was added to 3 drops of 10 μl of each of the fractions and the average number obtained represented the overall protozoa concentrations for each of the fractions. The prokaryotic community was obtained from the upper phase obtained during the protozoa sedimentation process and was centrifuged once at 500 ×*g* to remove potential remaining protozoa. Only the upper half of the supernatant was used to minimize contamination of protozoa after centrifugation.

### Microcosms preparation

The prokaryotic community was distributed evenly in 20ml anaerobic screw-cap glass tubes equilibrated in the anaerobic glove box. The rumen fluid containing the prokaryotic community was inoculated with 100mg of ground feed of the same composition the cows received as substrate. The protozoa fractions were centrifuged twice and concentrated in order to inoculate the microcosms with the smallest amount of volume to minimize the carryover of additional ruminal factors that might affect our experiment (150-250 μl, up to 2.5% of the final volume). The overall volume of each microcosm was 10 ml containing 10^4^ /ml of protozoa from each community, one treatment with the full native protozoa community (similarly adjusted to 10^4^ /ml cells) and one treatment without protozoa was named ‘protozoa-free’. The number of protozoa was chosen to reflect the typical abundance of protozoa in the rumen and was also based on a previous experiment showing that this number shows a visible change in methane production. The requirement for such protozoa numbers hindered our ability to always obtain the aimed triplicates for all the cows and fractions, thus some groups were performed with two replicates (cow 1; P<−10, cow2; All-protozoa, P-100). One additional all-protozoa community was removed from the analyses following as it showed contamination with a species of Betaproteobacteria, not native to the rumen. The microcosms were incubated for 96 h tilted at 20°. Methane quantification was performed after each day for four days. After methane quantification, 5ml of the upper fraction of the microcosms was removed and kept frozen at - 80□ for quantification of VFAs and sequencing of the prokaryotic community. The microcosm was complemented with 5 ml of medium M [20], and incubated further. All the procedures were performed under anaerobic conditions.

### Metabolites quantification

Methane and VFA quantification was performed following the protocol from Shabat et al (2016)[22]. For methane, the incubated samples were removed from incubation and directly placed into the Gas Chromatography (GC) autosampler 10 samples at a time. Samples of 0.250 ml of gas from the headspace of the tubes were injected into a 182.88 cm × 0.3175 cm × 2.1 mm packed Supelco analytical-45/60 Molecular sieve 5 A column (Supelco Inc., Bellefonte, PA, USA) with helium carrier gas set to a flow rate of 10 ml min–1 and an oven temperature of 200 °C. The oven temperature remained steady for a total run time of 5 min. A standard curve was generated using pure methane gas. After the daily measurement 5ml of fluid from each microcosm was removed for VFA quantification and microbiome analysis. For VFA measurement, the removed fluid was centrifuged at 10,000g in order to first separate the microbial community from the incubated fluid. The supernatant was transferred to a new tube and the pellet was used for further DNA extraction. 800 μl of the supernatant was mixed with 200 μl of 25% metaphosphoric acid solution (w/v in DDW) followed by 1 min vortex and then incubated at 4 °C for 30 min. The samples were then centrifuged for 15 min at 10,000 g and the supernatant was removed into new tubes, then 250 μl methyl tert-butyl ether (Sigma-Aldrich) was added and the tubes were vortexed for 30 s. Another cycle of centrifugation was performed for 1 min at 10,000 g. The upper phase, which contained methyl tert-butyl ether +SCFAs, was analyzed using an Agilent 7890B GC system (Agilent Technologies, Santa Clara, CA, USA) with a FID detector. The temperatures at the inlet and detector were 250 °C and 300 °C, respectively. Aliquots (1 μl) were injected with a split ratio of 1 : 11 into a 30 m × 0.32 mm × 0.25 μm ZEBRON ZB-FFAP column (Phenomenex, Torrance, CA, USA) with helium carrier gas set to a flow rate of 2.4 ml min–1 and initial oven temperature of 100 °C. The oven temperature was held constant at the initial temperature for 5 min, and thereafter increased at 10 °C min–1 to a final temperature 125 °C, and a final run time of 12.5 min. Individual injections of each pure VFA was performed in order to identify their retention in the column and a calibration curve was generated by preparing an equimolar solution of all the VFA and serially diluting it from 100mM to 0.1mM.

### DNA Extraction

DNA extraction was performed as previously described (Stevenson and Weimer, 2007). In brief, cells were lysed by bead disruption using Biospec Mini-Beadbeater-16 (Biospec, Bartlesville, OK, United States) at 3000 RPM for 3 min with phenol followed by phenol/chloroform DNA extraction. The final supernatant was precipitated with 0.6 volume of isopropanol and resuspended overnight in 50–100 μl TE buffer (10 mM Tris-HCl, 1 mM EDTA), then stored at 4°C for short-term use, or archived at −80°C.

### Illumina Amplicon Sequencing

The V4 region of 16S rRNA was amplified by PCR from DNA extracts using barcoded primers 515F 5′-CCTACGGGAGGCAGCAG-3’ and 806rcbR 5’-CCGTCAATTCMTTTRAGT-3’ [35]. The barcoded samples were pooled, sequenced in a MiSeq flow cell (Illumina) for 250 cycles from one end of the fragment and analyzed with Casava 1.8.

The Research Laboratory Hylab (Rehovot, Israel) performed amplicon sequencing for the 18S rRNA of the fractionated ruminal samples using primers specifically designed for ciliates taken from Tapio et al. (2016) [36] with the following sequences: CiliF (5’-CGATGGTAGTGTATTGGAC-3’) and CiliR (5’-GGAGCTGGAATTACCGC-3’). Ruminal DNA samples were treated as follows: 20 ng of DNA was used in a 25 μl PCR reaction with primers, using PrimeStar Max DNA Polymerase (Takara) for 20 cycles. The PCR reaction was purified using AmpureXP beads, and then a second PCR was performed using the Fluidigm Access Array primers for Illumina to add the adaptor and index sequences. For this reaction 2 μl of the first PCR were amplified in a 10 μl reaction for 10 cycles. The PCR reaction was purified using AmpureXP beads and the concentrations were measured by Qubit. The samples were pooled, run on a DNA D1000 screentape (Agilent) to check for correct size and for the absence of primer-dimers product. The pool was then sequenced on the Illumina MiSeq, using the MiSeq V2-500 cycles sequencing kit.

### Data analysis

Downstream processing of the 16S rDNA data, up to the generation of the ASV table was performed in QIIME v.2 [37]. DADA2 was applied to model and correct Illumina-sequencing amplicon errors and clustering of ASVs [38]. Taxonomic assignment for the bacterial 16S was performed using the pre-trained classifier Greengenes 13_8 99% ASVs from 515F/806R region from QIIME v.2 pipeline. After the generation of the ASV table, singletons/doubletons were removed and subsampling to an even depth of 4,000 reads per sample was performed for all subsequent analyses. Alpha and Beta diversity analyses were also performed using QIIME v.2 workflow, including principal coordinate analysis (PCOA) using the Bray-Curtis dissimilarity metric and ASV richness, evenness and Shannon diversity. Analysis of similarity (ANOSIM) was used to test the significance of the group clustering. For most statistical analysis of the compositional differences between the different microcosm groups, unless otherwise stated, ANOVA two-way performed on transformed data (using aligned rank transformation ART [39]), in order to assess the interaction between the different fraction groups and cow effect. When the analysis indicated a significant difference between the groups, a *post hoc* Aligned Rank Transform Contrasts (ART-C) was performed to determine which paired groups differed from each other [40] using the Artool package in r. When F values of ANOVAs on aligned responses not of interest did not meet the requirement of being ~0, as recommended by its developer [39], Kruskal-Wallis was performed along with the Wilcoxon test for pairwise comparisons between the groups.

For all the analyses, *P*-values of <0.05 after FDR correction were considered significant, unless otherwise stated in the text or figure. Statistical tests and data analysis across the different fractions were performed in R version 3.5.3 [41]. Multiple sequence alignment was performed using MAFFT, using the default parameters. The resulting multiple sequence alignment was used for the reconstruction of a maximum-likelihood phylogenetic tree using IQTree [42], with LG model and 1000 bootstrap replicates. The phylogenetic tree was visualized using iTOL [43].

## Results

### Experimental design

To study the effect of protozoa on the metabolic output and the prokaryotic community of the cow rumen, we performed a series of *ex-vivo* microcosm experiments. The experiments were initiated by sampling the rumen fluid of three cows that were kept under the same diet for two month prior to the experiment [20]. To produce protozoa communities characterized by different taxonomic composition we utilized the fact that different protozoa species have distinguishable sizes and shapes, and used an established approach in which the protozoa are fractionated by size [32, 44]. The protozoa community was fractionated into five fractions representing different protozoa sizes from >100μm to <10μm (Fig. 1a). Using this procedure we obtained protozoa communities that differ in taxonomic composition as characterized by 18s rRNA amplicon sequencing analysis (Fig. 1b). The fractions P-100 and P-60, were characterized by large protozoa mainly including *Ophryoscolex* and *Polyplastron* genera, with P-60 also including *Isotricha*. The P-40 fraction was almost exclusively composed of *Isotricha*. Fractions P-10 and P-<10 were dominated by *Dasytricha*, and to a lesser extent by *Entodinium*. The *Isotricha* genus was detected in all size fractions albeit in different relative abundances (e.g., between ~93% in P-40 to ~5% in P-<10) (Fig. 1b).

**Figure 1.**
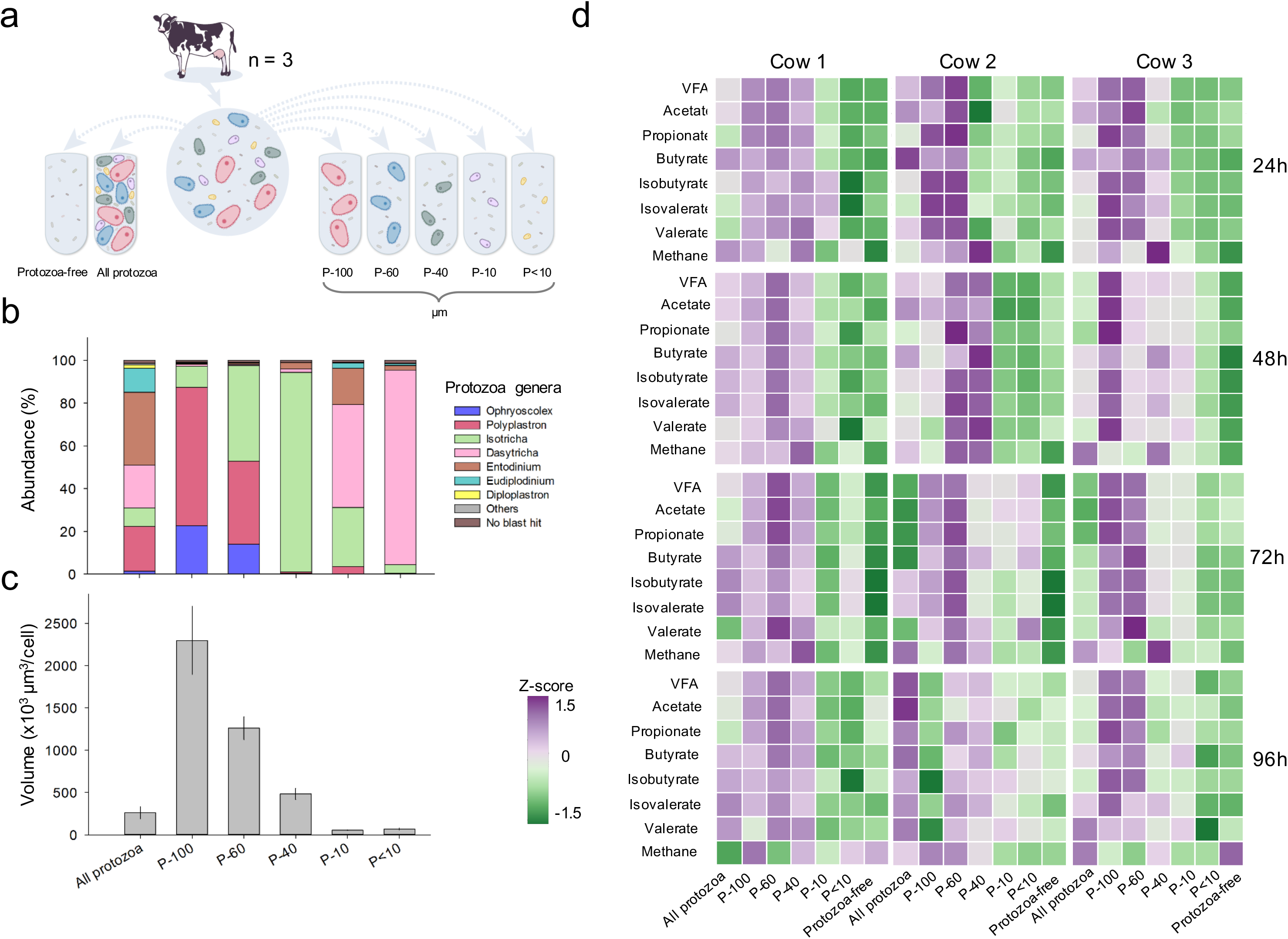
Experimental design and metabolic output of microcosms incubated with different protozoa communities. **(a)** Experimental setup of the microcosm experiments. The rumen microbial community of three cows were sampled and separated from protozoa cells. This was conducted either for all protozoa or according to the protozoa size indicated by P-100, P-60, P-40, P-10, P-<10 representing the different filters used for the separation. Subsequently, the microbial community was incubated with the different protozoa populations. **(b)** The genus level distribution of the protozoa in the different fractions obtained. **(c)** Average protozoa volume in each fraction obtained by measuring length and width of 25 cells per fraction. **(d)** Metabolic output of microbial communities incubated with different protozoa communities and without protozoa. Metabolites were measured every 24h in microbial communities from three cows (n=3 for each cow) for 96h that were incubated with different protozoa-size populations. Each row is represented by the z-score for each individual metabolite (for original data see Supplementary Fig. S1, Table S1).

After having established different protozoa communities, we set forward to measure their effect on an identical prokaryotic community. To this end, we adjusted the total number of protozoa cells for all subpopulations to 10^4^ cells/ml per tube (10^5^ cells overall in 10 ml medium) and exposed them to identical prokaryotic communities. Each protozoa community coming from each of the three cows were reintroduced to their respective native prokaryotic community of the cow they originated from. The resulting microcosms consisted of replicates of each communities along with replicate for the rumen prokaryotic community incubated without protozoa (hereafter referred to as ‘protozoa-free’ microcosms) and a fraction in which the whole protozoa population was reintroduced (hereafter referred to a ‘all-protozoa microcosms; Fig. 1a). Overall our experiment included three biological replicates per protozoa sub-population or control (21 experiments) and was conducted on three types of rumen bacterial communities originated from three different cows comprising an overall of 61 microcosms. The microcosms were incubated for 96 hours, and prokaryotic community composition as well as the methane and volatile fatty acid (VFA) production was assessed every 24 h.

### Methanogenesis is enriched in specific protozoal sub-communities

We first measured the effect of the different protozoal communities on ecosystem function, which was manifested by fermentation products and methanogenesis. Methane production is a classical example for a metabolic function that requires more than one microbial partner [45]. Our results show a clear and significant enrichment of methanogenesis in the protozoa fraction P-40 (Fig. 1d; FDR corrected Wilcoxon test *P* < 0.01). The P-40 community dominated by *Isotricha* exhibited a 1.5-fold higher methane output compared to fractions P-100, which exhibited the second highest methane production. Furthermore, P-40 protozoa exhibited a ~3-fold higher methane production compared to the protozoa-free microcosms after 48h (Fig. 1d). Overall, methane production in the small protozoa fractions P-10 and F<-10 was significantly lower than in the large protozoa fractions P-40, P-60 and P-100 (FDR corrected Wilcoxon test *P* < 0.05), but was still significantly higher when compared to the protozoa-free microcosms (Wilcoxon; *P* < 0.05; Fig. S1, Table S1). These results corroborate the role of protozoa in increased rumen methanogenesis and highlight the *Isotricha* dominated community as an high methane producing community.

Our results further reveal that the production of VFAs was significantly higher in several protozoa communities. Mainly, microcosms incubated with large sized P-100, P-60 and P-40 were significantly higher compared to the small P-10 and P-<10 protozoa communities and the protozoa-free community (Fig. 1d; Fig. S1, Table S1). Importantly, the effect of protozoa size on VFAs was maintained across all microcosms and time points (Fig. 1d). Linear regression based on size revealed that, after 48h, all VFAs, except valerate, exhibited a significant size dependence, with acetate exhibiting the strongest size dependence among the VFAs (Linear regression, R^2^ = 0.37 *P* = 10^−5^; Table S2). In contrast, methane production was only marginally dependent on protozoa size due the observed methanogenic enrichment in the intermediate size protozoa community P-40 (Fig. 1d; Linear regression, R^2^ = 0.24 *P* = 0.062; Table S2).

Our results show that the protozoa populations have a distinct effect on the overall metabolic output of the microbial community. The size of protozoa is highly linked to the effect on VFAs, yet, methane shows a pattern which suggests that additional factors may be in play.

### Protozoa sub-communities differentially shape prokaryotic community structure

To study the effect of protozoa on the microbial community structure in the rumen, we analysed the bacteria and archaea composition with relation to the different protozoa populations in the microcosm, across 96h of incubation via amplicon sequencing of the 16S rRNA in each microcosm. Our analysis revealed a clear and a strong causal effect of the distinct protozoa communities on the prokaryotic community structure in all of our biological replicates (microcosms and replicates coming from the different cows). Using the pairwise Bray-Curtis distance between the samples, a protozoa community based discrimination in prokaryotic community structure was already detectable after 24h and remained stable until the end of the experiment, as observed by the PCOA clustering of the prokaryotic community as a function of the different protozoa populations (Fig. 2a; Fig. S2). Furthermore, replicates inoculated with the same protozoa community were significantly more similar to each other than between the different protozoa communities (*P* = 1e-9 using Wilcoxon test; Fig. 2a). The individual effect of the protozoa communities on prokaryotic structure was evident in all three rumen prokaryotic communities originating from the different cows, despite the large differences stemming from the individual cows (Fig. S3; ANOSIM R= 0.98, *P* < 0.0001), and dynamics across the days of sampling (Fig. S3). Interestingly, the time dependent difference between 72h and 96h was marginal, while the differences in community structure stemming from the different protozoa communities remained stable (Fig. S3). To further evaluate the strengths of change in prokaryotic community structure induced by the different protozoa communities, we compared the Bray-Curtis distance between protozoa-free to protozoa-containing communities, which revealed that Bray-Curtis distance was largely dependent on the size of the protozoa (Fig. 2b). Large protozoa cells (P-100, P-60 and P-40) exhibited a higher distance than smaller protozoa sizes (P-10 and P-<10) compared to the protozoa-free microcosms (Fig. 2c). Interestingly, the largest distance from the protozoa-free population was observed in the fraction containing the native protozoa population despite encompassing a lower average protozoa volume (i.e., all-protozoa fraction, Fig. 2c).

**Figure 2.**
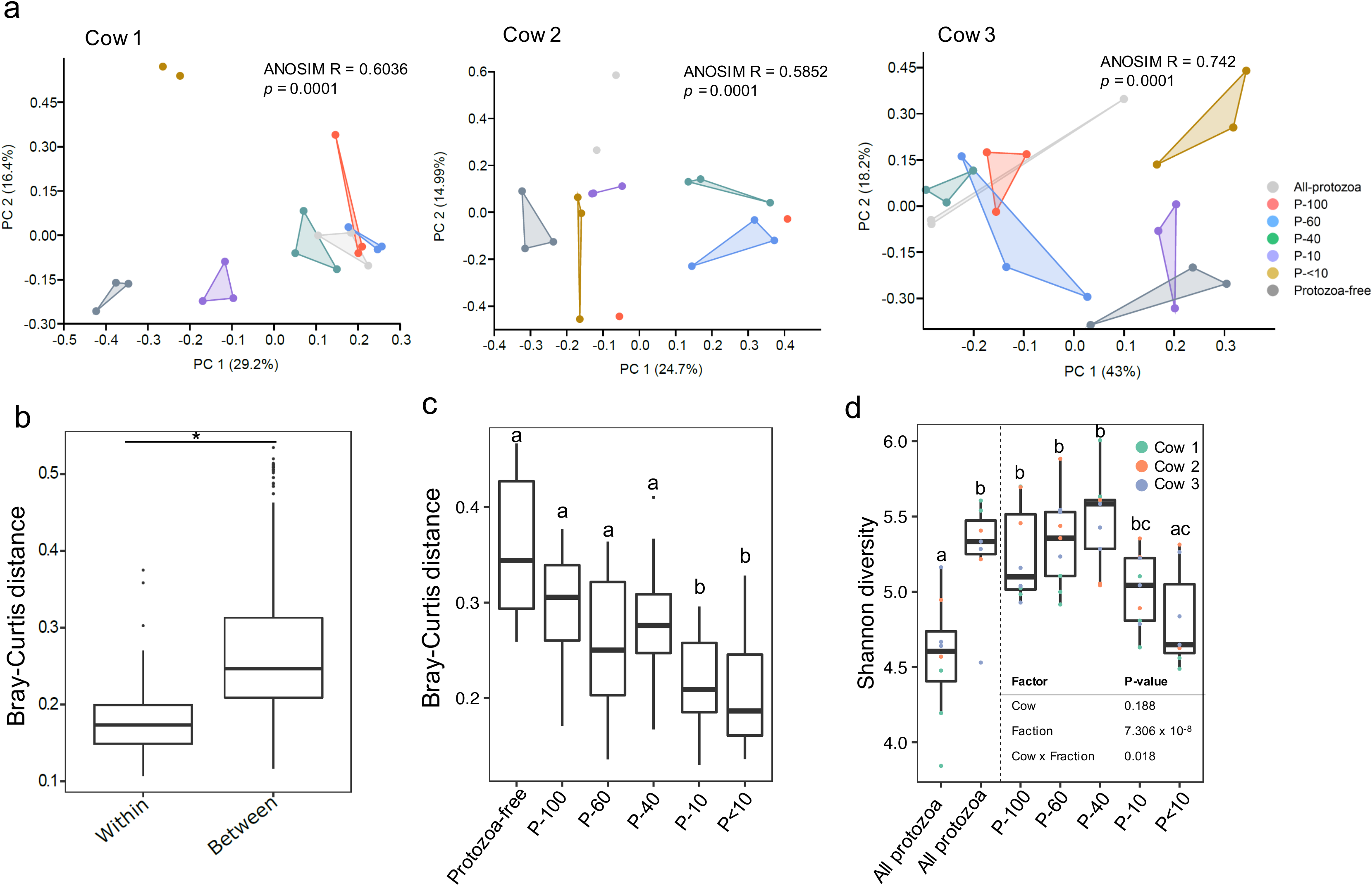
Ecological structure of the prokaryotic community across microcosms. **(a)** Principal coordinate analysis (PCOA) plot of the microcosms separately plotted according to the cow the microbiome originates from (cow #1-3) and based on Bray-curtis distance metric at the end of the experiment. Analysis for all time points of the experiment are in the ‘supplementary material’ (Fig. S2). Analysis of similarity (ANOSIM) was performed to assess the protozoa community-mediated discrimination (shown in upper right corner of each plot). **(b)** Bray-Curtis distance within replicates across the microcosms compared to the distance between the replicates. The values used are the individual values obtained for each source (i.e., cow) sample. Pairwise Wilcoxon test was used to test for significance (*P* = 10^−9^) **(c)** Bray-curtis distance between the protozoa-free microcosms and microcosms containing different protozoa communities. Pairwise Wilcoxon rank sum test with FDR correction was used to test for significance; boxes that are not sharing a letter denote significance at *P* < 0.05. **(d)** Shannon diversity across the different treatments at the end of the experiment. The table on the lower right side of the plot shows the results of ANOVA two-way performed on transformed data (using aligned ranked transformation [39]) to test the overall difference between the groups and interaction between parameters (cow effect * protozoa community effect). A *post-hoc* pairwise FDR corrected test was performed for significance between the groups with boxes that are not sharing a letter denoting a significant difference between the groups at *P* < 0.05.

We further analysed the effect of the inoculated protozoa communities on the alpha diversity parameters of the prokaryotic communities. While we did not observe a consistent difference in diversity at the first two days of incubation (Fig. S4), over time (after 72 h and 96 h) microcosms incubated with P-100, P-60 and P-40 invariably resulted in a significantly higher Shannon diversity, species richness and evenness than the protozoa-free community (Aligned Rank Transform (ART) ANOVA *P* < 0.01; Fig. 2d; Fig. S4). Notably, communities incubated with the smaller P-10, P-<10 protozoa showed a trend in increased diversity (Fig. 2d). Interestingly, the fraction P-40, representing an intermediate protozoa size, exhibited the highest prokaryotic diversity in most microcosms compared to the protozoa-free microcosm (Shannon diversity; Protozoa-free = 4.58, P-40 = 5.47; *P* < 0.001, FDR corrected ART-C test), driven by a higher ASV richness (Fig. S4).

Our results show that the presence of protozoa populations had a strong impact on the rumen microbial ecosystem diversity and that different protozoa communities differentially affect prokaryotic community structure. Protozoa of larger size and volume induced stark alterations in the microbial community structure, while prokaryotic community richness peaked in the intermediate sized protozoa fractions.

### Protozoa positively affect specific rumen prokaryotic lineages with a strong effect on the enrichment of Gammaproteobacteria

To quantify the effect of protozoa on the abundance of specific prokaryotic species, we analysed the taxonomic distribution across all samples. The analysis yielded 14 classes, 26 families, and 31 genera that were present on average above 0.5% of the total prokaryotic community in at least one group of replicates and represented between 85% to 97% of the total prokaryotic community. We observed a large expansion of Proteobacteria in all populations, chiefly attributed to Gammaproteobacteria, already observable after 24h (Fig. 3a; Fig. S5). The increase in Gammaproteobacteria abundance was most pronounced in communities incubated with large protozoa fractions (P-100, P-60, P-40), where the increase ranged between 3-fold to 20-fold higher bacterial abundance (Wilcoxon test, *P* < 0.05; Fig. 3a; Fig.S5). Notably, one class of methanogenic archaea, Methanobacteria, exhibited a clear higher proportion only in fractions P-100, P-60 and P-40, with the latter fraction exhibiting the highest relative abundance (Fig 3a, Fig. S5). Bacteroidia, the most abundant class in the samples, exhibited a trend of lower abundance in protozoa-containing microcosms, but was overall highly variable between cows and replicates (Fig. 3a; Fig. S5).

**Figure 3.**
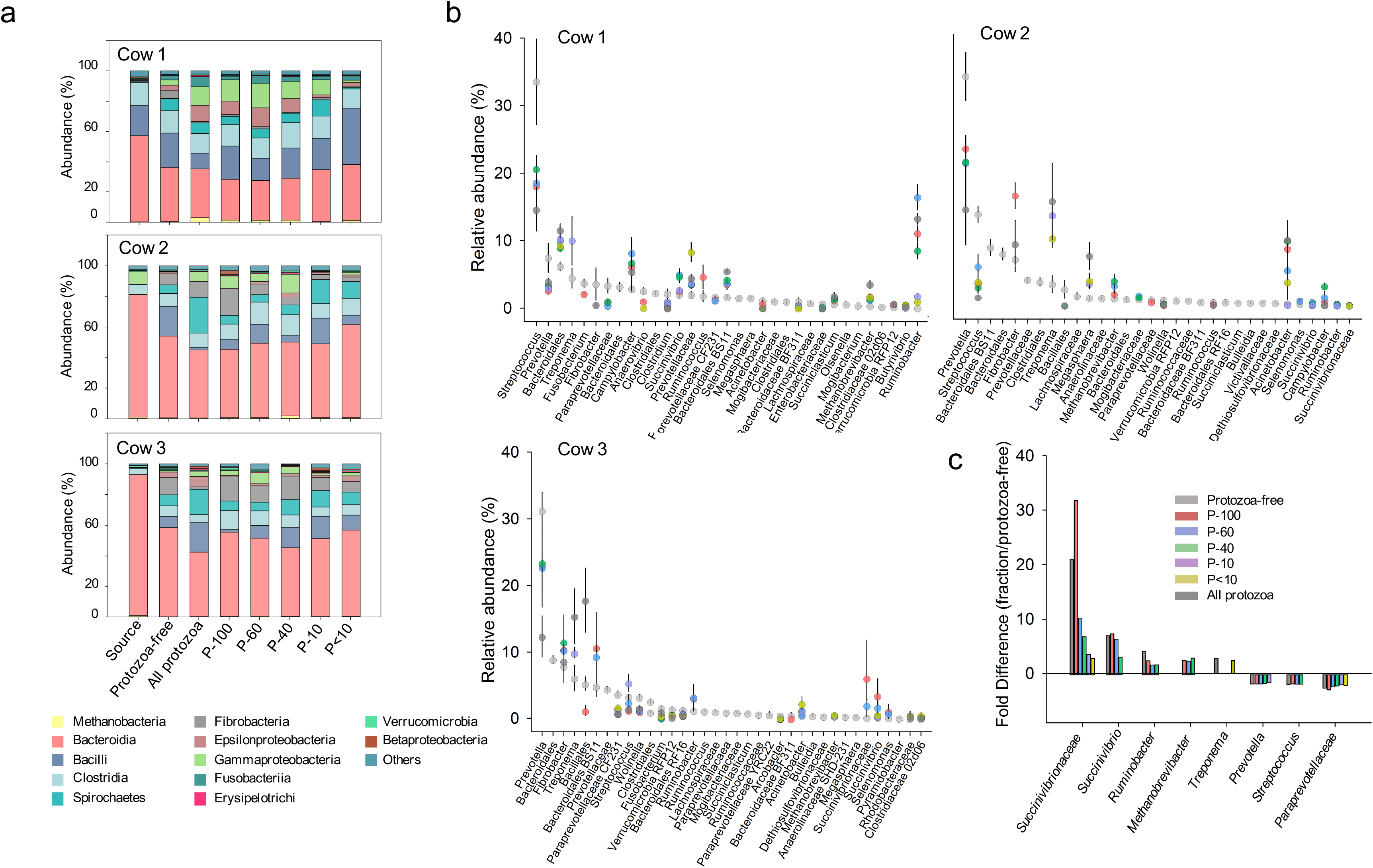
Taxonomic composition across the microcosms. **(a)** Stack bars displaying the class level relative abundance of taxa across the different protozoa communities, the protozoa-free community and for each cow, the original prokaryotic composition of the samples taken (termed ‘source’). **(b)** The relative abundance of the different genera in the microbial communities is displayed for each cow. The genera are ordered based on their rank abundance in the protozoa-free microcosms. Color coding is based on the different protozoa communities added to the prokaryotic community and only genera that were significantly different from the protozoa-free fraction are displayed (FDR corrected t-test P < 0.05) (d) Summary fold-differences of taxa that exhibited the same directional variance across the different source (i.e. cow) between the protozoa-free microcosms and the communities containing different protozoa communities. Color coding is based on the different fractions.

To further assess taxa distribution, we conducted a genus level analysis that showed that, depending on the source of the prokaryotic community (i.e., cow), bacterial genera enriched within the Gammaproteobacteria were Succinivibrionaceae, *Succinivibrio, Ruminobacter, Acinetobacter* or all together (Fig. 3b,c). These genera represented on average between 1.4% to 30% in the protozoa-containing communities representing between 2 - 20 average fold increase compared to the protozoa-free microcosms (Fig. 3b,c). These genera were in either lower abundance or completely absent in the protozoa-free microcosms ranging between 0% to 4.7%. The increase in Gammaproteobacteria was contrasted mainly by *Prevotella* and *Streptococcus,* which were usually the most abundant genera in the microcosms and exhibited a significant decrease across all cows (Fig. 3b,c).

Our results show that the presence of protozoa had a recurrent effect on the bacterial composition, regardless of its source community, particularly with regards to the enrichment of specific taxonomic lineages (Fig. 3c).

### Protozoa favour co-existence of phylogenetically related taxa

Our results of the prokaryotic taxonomic distribution so far show that protozoa had a stark effect on the microbial community composition. To assess the protozoa-mediated effect on the microbial species level, we conducted a phylogenetic analysis of ASV-level taxa (Fig. 4a). This revealed that ASV-level taxa largely reflected the taxa distribution observed at the genus level (Fig. 4a). Interestingly, we observed a significantly larger proportion of ASVs that increased in abundance in the presence of protozoa were shared between the different cows compared to ASVs that decreased (Fig. 4a,b). Only 6 out of 40 ASVs significantly decreased in protozoa-containing microcosms, which was consistent across at least two cows (3 ASVs shared by all three cows). In contrast, the number of ASVs exhibiting a higher abundance in protozoa-containing microcosms was significantly higher with 39 ASVs shared between cows (Fisher exact test increasing ASV vs. decreasing ASV; *P* < 0.001; Fig 4a,b). Our results thus show that taxa enriched in the presence of protozoa were more consistent across the different communities originating from different cows, suggesting that protozoa confer an advantage to specific prokaryotic lineages. In contrast, ASVs that exhibited a decreased presence were significantly less recurrent across the different cows, thus likely more dependent on the initial composition of the microbial community (Fig. 4b).

**Figure 4.**
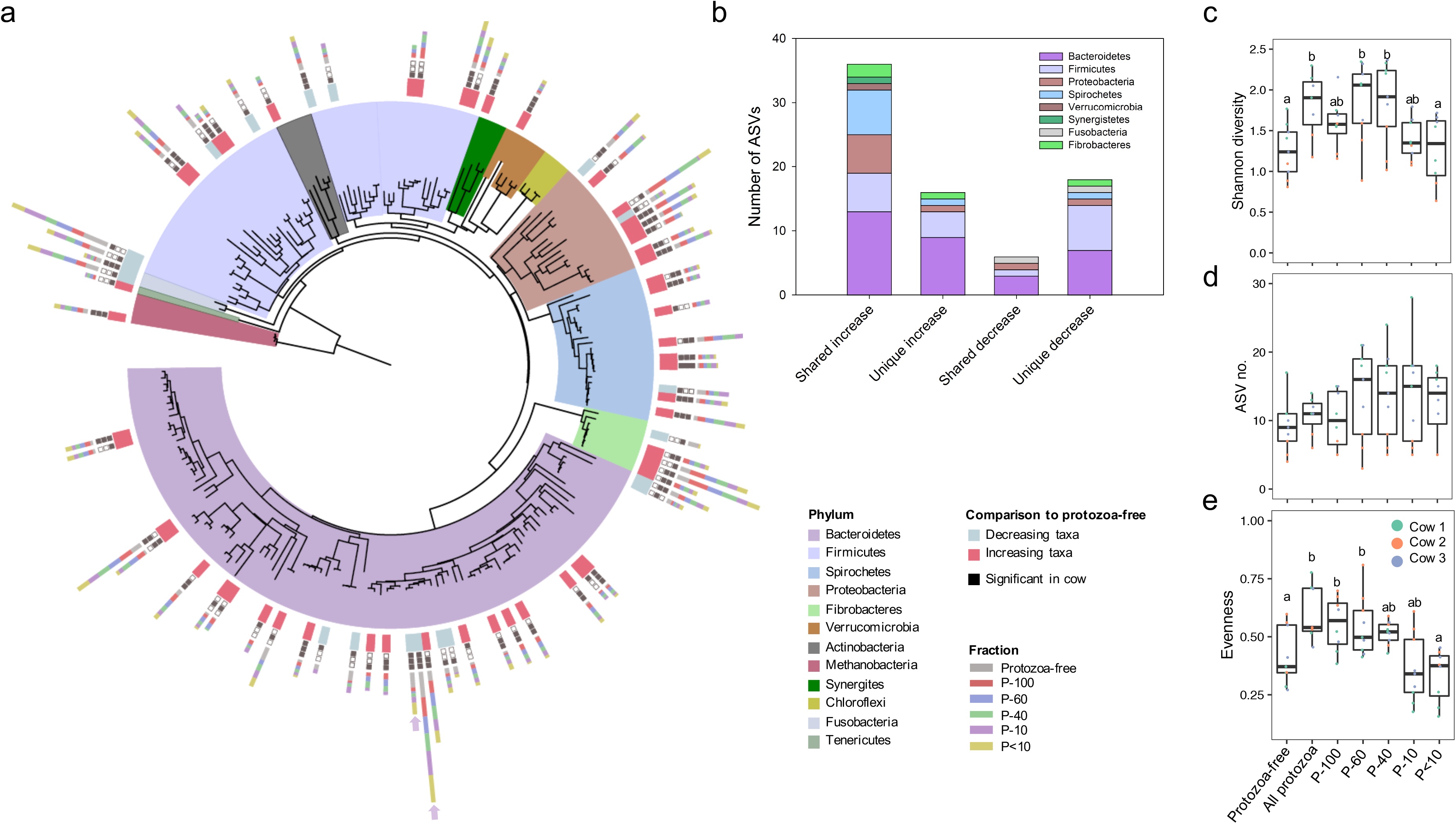
ASV distribution across the microcosms. **(a)** Phylogenetic tree of the ASVs that were above 0.5% relative abundance in at least one group of microcosms. Each ASV is color coded based on their phylum affiliation. The colored boxes above each ASV represent their divergence in abundance in the protozoa-containing microcosms compared to the protozoa-free microcosms (red=increasing, light blue=decreasing). The filled and empty black squares represent the cows in which the difference was observed with the filled square denoting that a difference was observed in a specific cow. The stack bars represent the average abundance of the ASV across the different communities. **(b)** Distribution of differentially abundant ASVs in protozoa-containing microcosms compared to the protozoa-free microcosms. The stack bars denote whether the ASVs exhibited a significant increase or decrease and whether they were shared across cows or unique to one cow. **(c)** Shannon diversity **(d)** no. of ASVs and **(e)** evenness of the genus *Prevotella* across the different communities. (**c-e)** Differences between the groups was assessed using aligned rank transformed ANOVA *post hoc* test ART-C, with different letters above the boxes signifying significant differences between the groups at *P* <0.05.

Further analysis of the distribution of taxa that are differentially abundant when incubated with protozoa, we observed that they include a large proportion of *Prevotella* and *Treponema* associated ASVs, which are either significantly higher in abundance or exclusively found in protozoa-containing microcosms (Fig. 4a,b). Interestingly, *Prevotella* overall exhibited a decrease in abundance in most of the protozoa-containing microcosms (Fig. 3). We found that while overall decreasing in abundance, *Prevotella* diversity significantly increased in the presence of protozoa (protozoa communities P-60, P-40 and all-protozoa, *P* < 0.05 FDR corrected Aligned Rank Transform Contrast (ART-C) ANOVA test; Fig. 4c-e). The increase in diversity was mostly driven by an increase in ASV richness and evenness within the genus depending on the source of the prokaryotic community (Fig. 4e), Concomitant with a decrease of the dominant *prevotella* ASVs found in the protozoa-free microcosms (Fig. 4a, arrows). An even more striking observation could be made within the *Treponema* genus (Fig. S6). This genus overall increased in abundance across time in all the microcosms regardless of whether these contained a protozoa community or not (Fig. S6). However, the expansion of *Treponema* in protozoa-free microcosms was limited to a small number of ASVs, while it accounted for significantly more ASVs in the all protozoa-containing microcosms (*P* < 0.05 FDR corrected ART-C test; Fig. 4a,b; Fig. S6). These results thus show that the presence of protozoa is directly responsible for the increase within genus diversity. Thus, in addition to the selection of specific prokaryotic lineages, the presence of protozoa may also allow for an increased co-existence of phylogenetically similar taxa within microbial communities.

## Discussion

Host-associated microbial communities can be altered by several factors including the host species or genetics [26, 46], host lifestyle such as diet, and geography of the host [47]. In addition, within the constraint of these parameters, microbiome internal parameters, such as interactions between its species, can shape community structure as well [48, 49]. While bacteria-bacteria interactions have garnered considerable attention in recent studies, the effect of the eukaryotic components of microbial communities remains largely unexplored. Here we establish an experimental system that enables us to control for the presence and absence, as well as the composition of the protozoa community. Our experimental set-up allows us to show that the presence of different protozoa communities leads to significant individual changes in prokaryotic community structure as well as end-product metabolite output including methane (Fig. 5).

**Figure 5.**
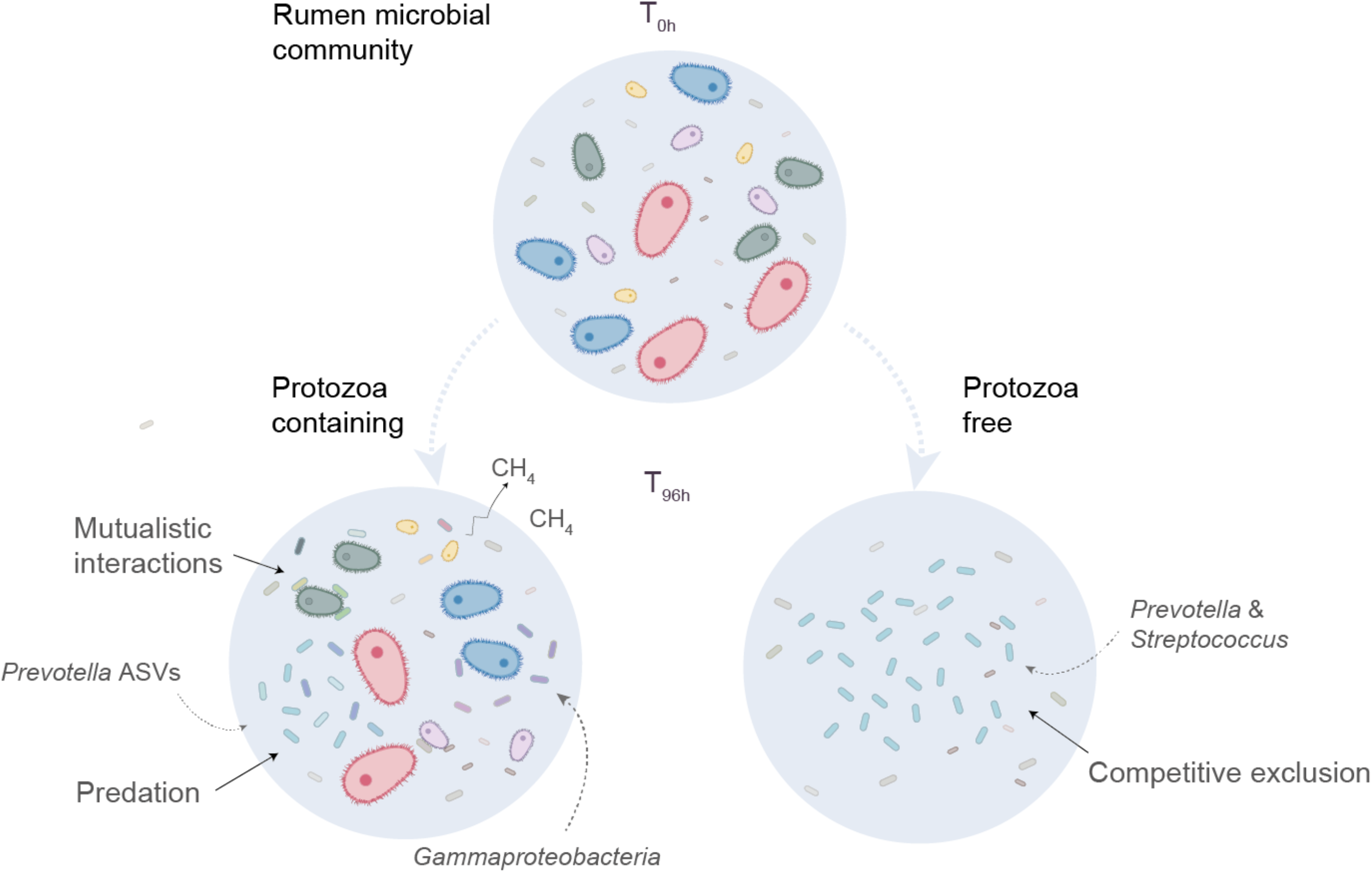
Modulatory effect of protozoa on the rumen prokaryotic community as seen in this experiment. The microbiome samples were originally taken from a mixed community of bacteria, archaea and protozoa. When removed from the microbial community, prokaryotic diversity was lower than when protozoa remained in the system. This suggests that protozoa maintain diversity of the prokaryotic community of the rumen and enable the co-existence of phylogenetically related species (represented by the different shades of blue denoting *Prevotella* species). The presence of protozoa also enriches specific bacterial lineages such as Gammaproteobacteria either by metabolic mutualistic interaction or by resistance to predation of this taxon.

The protozoa-mediated enrichment of specific taxa, such as the expansion of Gammaproteobacteria families and genera, suggests metabolic interactions between the bacteria and protozoa. Indeed, species of *Succinivibrionaceae* have been characterized as hydrogen utilizers, which protozoa are known to produce in high abundance via their hydrogenosomes [17, 50]. This observation can be extended to the concurrent increase in *Methanobrevibacter*, a methanogenic archaea genus, in which most of its known species are hydrogen utilizers (Fig. 3). Nonetheless, the rumen environment encompasses a large diversity of taxa capable hydrogen utilization, with a recent study analyzing 501 rumen genomes showing that two third of those carry hydrogen utilization/production capabilities [51]. The expansion of Gammaproteobacteria may thus be the result of additional factors that likely confer an advantage over competing species in the presence of protozoa. Interestingly, Gammaproteobacteria were previously suggested to be resistant to predation in marine microbial assemblages [52, 53]. It was further suggested that types III, IV and VI secretion systems, which are commonly encoded in the genome of this class, may assist in the observed resistance to predation [52, 54]. Notably, type III secretion systems were also identified as highly abundant in Gammaproteobacteria species in the rumen environment [55]. In addition, Gammaproteobacteria genera in the rumen environment were shown to display a large variability in abundance between animals, even under similar management and diet [56, 57]. Notably, *Succinivibrionaceae* were observed to be associated with higher feed efficiency and lower methane emissions in ruminants and other foregut hosts [58, 59]. Therefore, our results may offer an explanation to such variations, where protozoa composition and abundance play a role in enriching Gammaproteobacteria, subsequent metabolic output and animal phenotype.

In their role as microbial predators, a large body of theoretical framework as well as empirical evidence show that protozoa modulate the relationship between microorganisms by exerting top down control on the overall structure of prey communities [7, 60–62]. Predators are often considered keystone species in an ecosystem, as they are able to impose a strong selection on prey communities even when they are found in low abundance. Bactivorous predation by protozoa is often considered of a generalist nature, where a wide breadth of feeding preference can be observed largely affecting bacterial density, but less so the overall community composition [5]. This is in contrast to selective predation, which has the potential to extinguish entire clonal populations, thus changing prokaryotic composition. In the rumen, Gutierrez (1958) observed that *Isotricha prostoma* preferentially ingested bacteria of specific morphologies [63]. In contrast Coleman (1964), observed that *Entodinium caudatum* had no preference in bacterial prey when offered bacterial mixtures with differing proportions [64]. Based on our observation of a stark change in community composition, it is likely that selectivity in prey species exists in the rumen protozoa populations. One reason for such observation may be adaptation of prey species via increased resistance to predation.

Microcosm experiments as well as theoretical models previously demonstrated that exposure to predation offsets competition between species with overlapping niches in a competition-predation trade-off [60, 65, 66] ultimately leading to an increase in diversity parameters. This is in line with our findings, where the presence of protozoa in the rumen community increases diversity that is driven by both increased evenness and ASV number by the end of the experiment (Fig. 2). Our results further strengthen this notion, as we find that the presence of protozoa decreases the abundance of the most dominant taxa such as *Prevotella* or *Treponema*, concomitant with an increase in diversity of phylogenetically related taxa (Fig. 4). These genera are an example of the protozoa-mediated effect in mitigating competition between phylogenetically and possibly metabolically similar taxa. Our results thus further suggest that protozoa, potentially via predation, play a central role in promoting species coexistence of species with overlapping niches in complex microbial communities. Such a role of promoting diversity in host-associated communities was also proposed for the eukaryotic community in the human gut [15, 67, 68].

Interestingly, the increase in diversity parameters was most pronounced in fractions containing intermediate sized protozoa (P-40, Fig. 2). This observation may be interpreted as a result of a competition-predation trade-off, where protozoa-free microcosms and microcosms containing only large protozoa (P-100) represent two extreme scenarios leading to taxa extinction due to either high competition (protozoa-free) or strong predation (large protozoa). In contrast, intermediate sized protozoa may represent an equilibrium between the competition-predation trade-offs, displaying a high species diversity. This scenario fits prior experimental models, which show that prey diversity is maximized at intermediate predation intensity [69, 70]. However, validating this hypothesis requires further experimentation that would include a decoupling of protozoa size from their identity.

Our results further show that protozoa play a pivotal role in the rumen microbiome end-product output comprising VFAs and methane (Fig. 1). While the higher production of several of the quantified metabolites such as acetate and butyrate may be related to the protozoa metabolism [29], methane is likely the result of mutualistic interactions between methanogenic species and hydrogen producing microbes. Several microbial eukaryotes form mutualistic (or commensal) relationships with prokaryotes across a wide range of environments [2, 71]. Indeed, rumen protozoa were shown to be habitat for a large methanogenic community that is physically associated with the protozoa cells [32, 34, 72]. Consequently, the elevated methane emission was suggested to be the result of a mutualistic relationship between the hydrogen producing protozoa and the hydrogenotrophic methanogens. Our results are in line with this observation and further show that the hydrogen-utilizing methanogenic community increased in the presence of larger protozoa species (Fig. 3). Thus, the strong increase in methane emission measured in the presence of protozoa (especially P-40), is likely explained by the protozoa-associated microbial community.

Many experiments studying the effect of micro-eukaryotic predators on the bacterial community use artificial prey communities that comprise only a low number of different species or a priorly simplified bacterial community [5, 60, 73]. Here, we assessed the direct impact of the presence and absence of natural protozoa communities on their native prokaryotic communities. Accordingly, our study provides insights into natural dynamics as well as the multifaceted role of microbial eukaryotes in microbial habitats. Protozoa feed on the microbial populations, yet they also provide habitats and nutrients for mutualistic exchange to their surrounding prokaryotes. Thus, when studying the rumen microbial ecosystem, the role of cross-domain interactions between protozoa and prokaryotes need to be taken into consideration. Modulation of ciliates may bear great potential in affecting its surrounding prokaryotic community, which may also lead towards improved animal phenotypes.

## Acknowledgments

Our gratitude goes to the ARO farmer and veterinarians for their support throughout this experiment. We want to thank Ido Toyber for his critical reading of the manuscript. We thank Fenna T. Stücker for the graphical design of the experimental scheme.

## Funding

This study was supported by grants from the Israeli Dairy Board foundation (Grant No. 362-0524/25) and the Israeli Science Foundation (Grant No. 603/20).

## Data availability

The dataset analyzed during the current study will be promptly available in the NCBI Sequence Read Archive (SRA) database.

## Author information

### Contributions

EJ and BL conceived the experiment. EJ, RS, BL, VR, RD, SE performed the incubation experiment, metabolite quantification, DNA extraction and sequencing. RS, TZ and OF performed the sequencing. EJ, RS, IM and TW analysed the data. EJ, TW, IM and RS wrote the manuscript. All authors read, revised, and approved the final manuscript.

## Ethics declarations

### Consent for publication

Not applicable.

### Conflict of interest

The authors declare that they have no conflict of interest.

## Notes

### Competing Interest Statement

The authors have declared no competing interest.

