## Supplementary Material for "Protozoa populations are ecosystem engineers that shape prokaryotic community structure and function of the rumen microbial ecosystem"

### Supplementary Figures

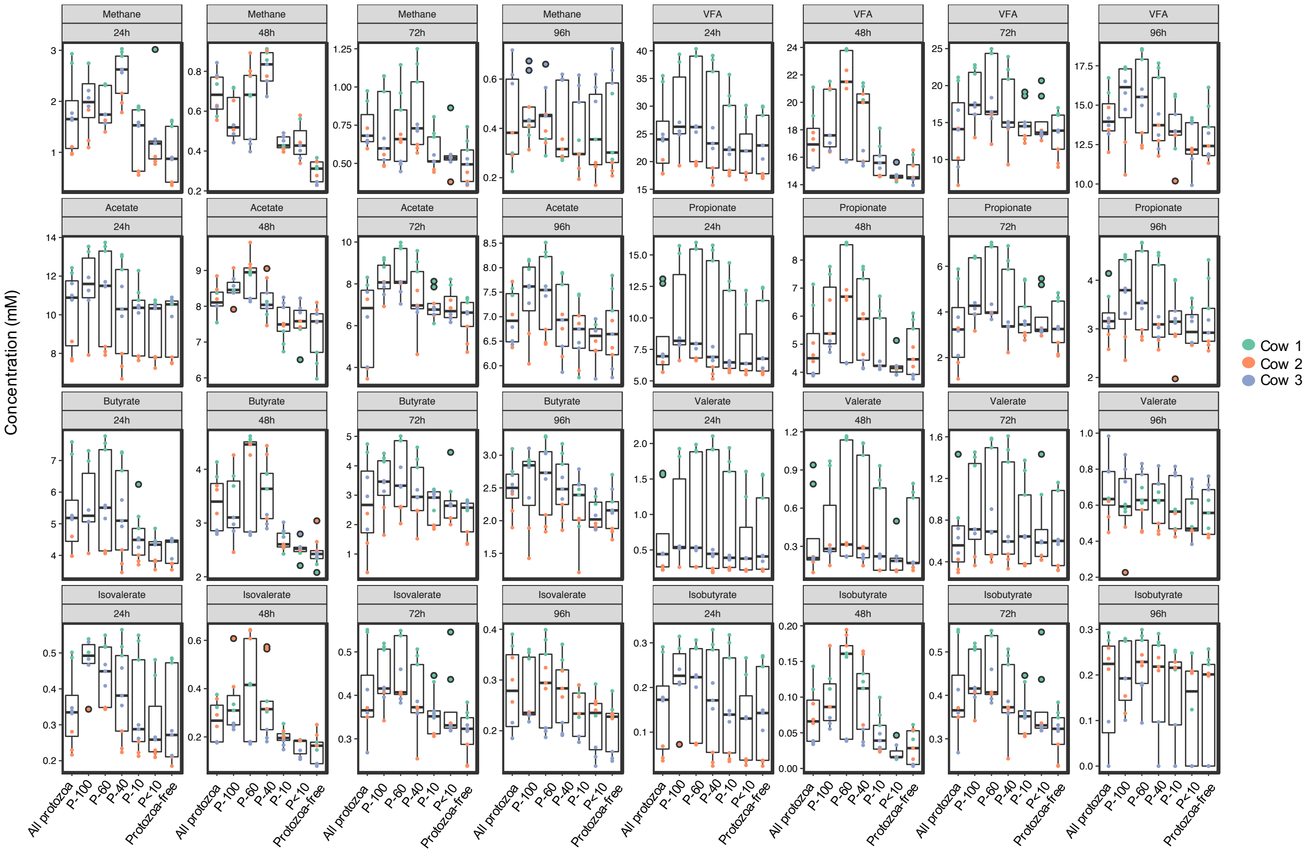


**Supplementary Figure S1. Production of VFAs and methane over time.** Metabolites were measured in microcosms containing the rumen microbial community incubated with different protozoa communities and protozoa-free microcosms. VFAs and methane were measured every 24h for a total of 96h. Kruskal-Wallis statistical test was performed to test for differences between the protozoa fractions and are shown in Table S1.

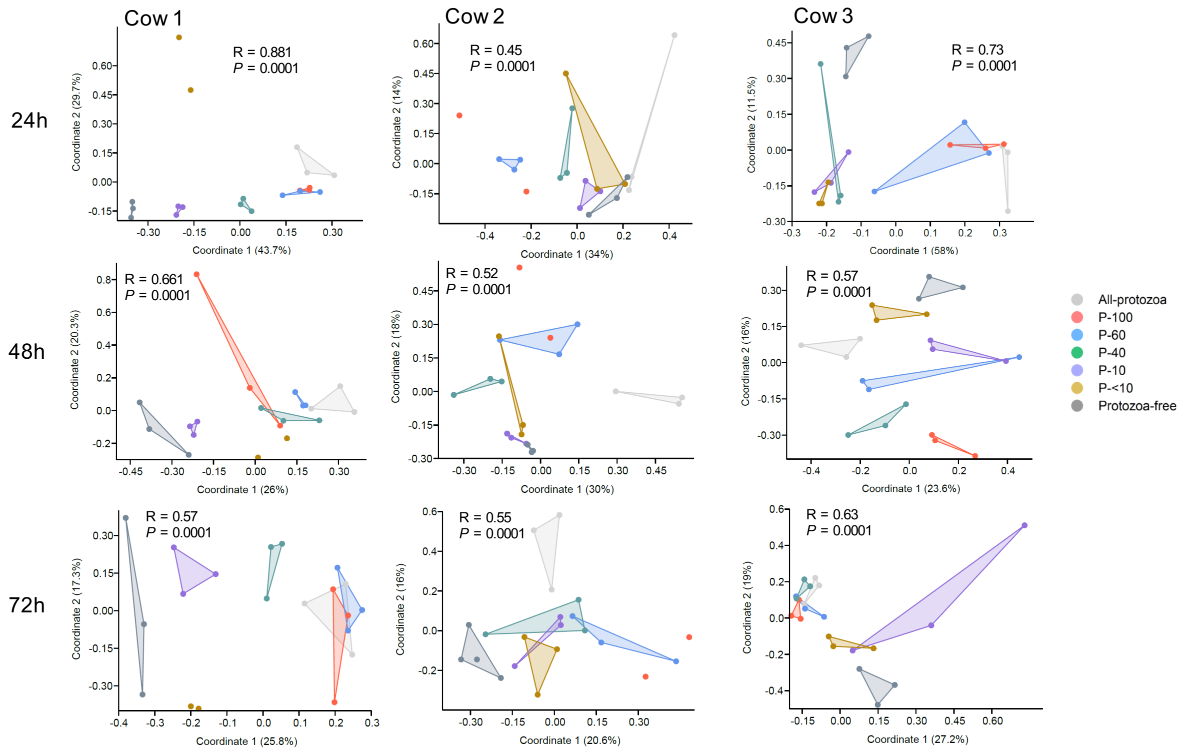


**Supplementary Figure S2. Ecological structure of the prokaryotic community across time.** Principal coordinate analysis (PCOA) based on pairwise Bray-Curtis distance metric of the microcosms plotted by source sample and for every day of the experiment (24h to 72h; 96h is shown in Fig. 2a). Analysis of similarity test (ANOSIM) on the overall groups was performed and displayed for each plot. This analysis shows that discrimination based on the different protozoa communities introduced to the prokaryotic communities, can be detected after 24h.

a

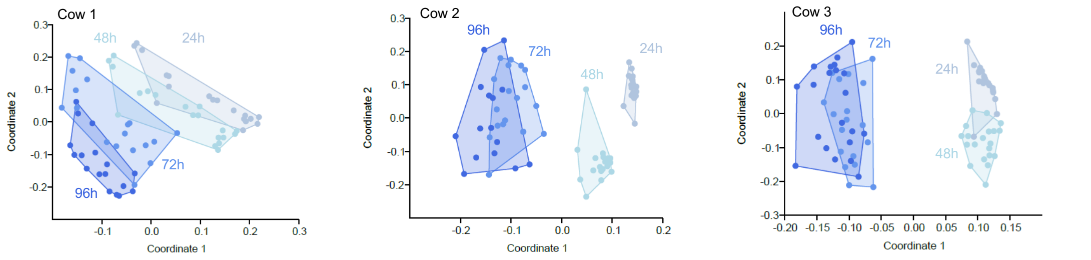


| R/p-value | 24h | 48h | 72h | 96h |
|---|---|---|---|---|
| **24h** | | 0.001 | 0.001 | 0.001 |
| **48h** | 0.2215 | | 0.001 | 0.001 |
| **72h** | 0.6249 | 0.434 | | 0.04 |
| **96h** | 0.8864 | 0.7192 | 0.343 | |

| R/p-value | 24h | 48h | 72h | 96h |
|---|---|---|---|---|
| **24h** | | 0.001 | 0.001 | 0.001 |
| **48h** | 0.8636 | | 0.001 | 0.001 |
| **72h** | 0.9748 | 0.9014 | | 0.046 |
| **96h** | 0.9913 | 0.9375 | 0.1522 | |

| R/p-value | 24h | 48h | 72h | 96h |
|---|---|---|---|---|
| **24h** | | 0.001 | 0.001 | 0.001 |
| **48h** | 0.7253 | | 0.001 | 0.001 |
| **72h** | 0.973 | 0.9686 | | 0.06 |
| **96h** | 0.9705 | 0.9586 | 0.1604 | |

b

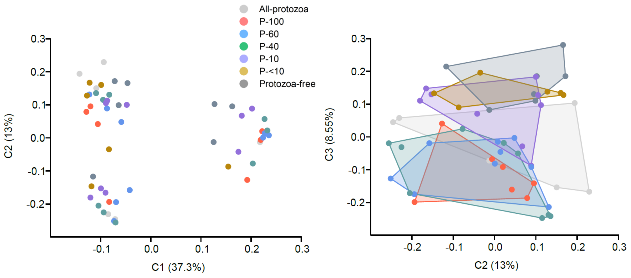


| R/p-value | All-protozoa | P-100 | P-60 | P-40 | P-10 | P-<10 | Protozoa-free |
|---|---|---|---|---|---|---|---|
| **All-protozoa** | | 0.2512 | 0.1333 | 0.3179 | 0.0356 | 0.0341 | 0.0041 |
| **P-100** | 1.263 | | 0.9382 | 0.9207 | 0.0948 | 0.03 | 0.0197 |
| **P-60** | 1.712 | 0.3458 | | 0.8035 | 0.0958 | 0.0092 | 0.0123 |
| **P-40** | 1.152 | 0.3057 | 0.4547 | | 0.0951 | 0.0058 | 0.0044 |
| **P-10** | 2.35 | 1.786 | 1.989 | 1.872 | | 0.1459 | 0.2584 |
| **P-<10** | 2.829 | 2.984 | 3.988 | 3.836 | 1.804 | | 0.163 |
| **Protozoa-free** | 3.367 | 2.618 | 3.116 | 3.308 | 1.253 | 1.619 | |

**Supplementary Figure S3. Effect of incubation time and cow individuality on microbiome structure.** (a) Principal coordinate analysis (PCOA) based on pairwise Bray-Curtis distance metric of the microcosms plotted for the different cows and colored based on the day of sampling of the microcosms after the beginning of incubation (every day from 24h - 96h). The table below each plot denotes the ANOSIM R values and their corresponding P-values between the different days. We can observe rapid dynamic change between the first 3 days which subsides on the 4th day, and marginally different when compared to the third day. (b) Effect of cow individuality on prokaryotic structure. Left, the PCOA shows 3 separate clusters representing each cow and color coded based on the protozoa community present in each sample at the end of the experiment. This demonstrates the strong effect of the origin of the sample on prokaryotic composition. Right, PC2 and PC3 of the PCOA showing a discrimination based on the protozoa communities, demonstrating that features of the prokaryotic community are dependent on the protozoa communities added to the prokaryotic community and shared between the cows. The table shows a PERMANOVA analysis across the protozoa communities.

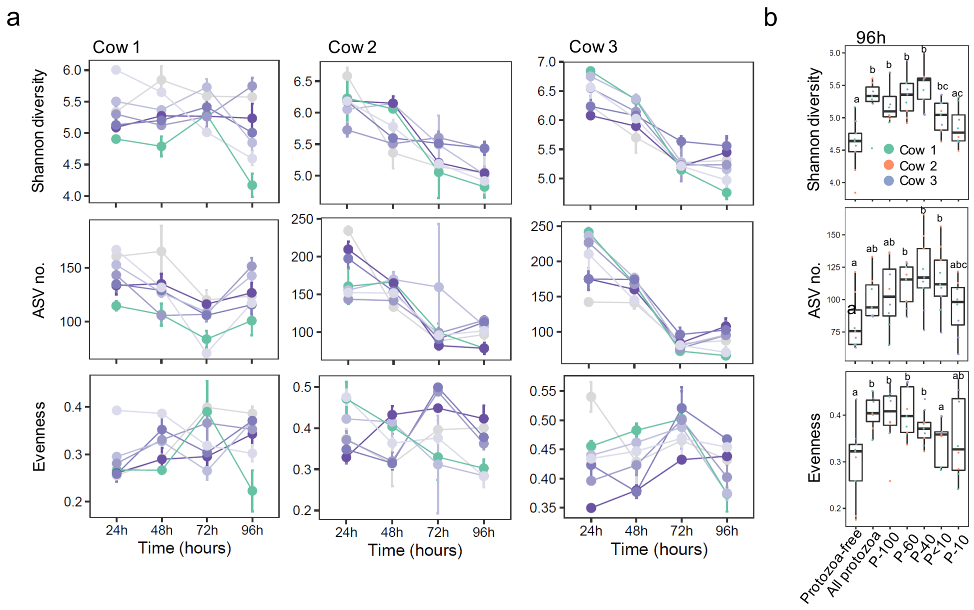


| Factor | P-value |
|---|---|
| Cow | 0.0445380 |
| Protozoa fraction | $7.3069 \times 10^{-7}$ |
| Cow x Protozoa fraction | 0.0092896 |

| Factor | P-value |
|---|---|
| Cow | $6.882 \times 10^{-7}$ |
| Protozoa fraction | 0.0004294 |
| Cow x Protozoa fraction | 0.2063852 |

| Factor | P-value |
|---|---|
| Cow | $2.4577 \times 10^{-7}$ |
| Protozoa fraction | $2.0210 \times 10^{-6}$ |
| Cow x Protozoa fraction | 0.011215 |

**Supplementary Figure S4. Prokaryotic alpha diversity across all microcosms.** (a) Alpha diversity parameters plotted for individual cows and across time from 24h to 96h including ASV richness (ASV no.), Shannon diversity and evenness across the different microcosm groups. We observed different dynamics across the different cows which invariably lead to higher diversity parameters in microcosms containing protozoa of larger sizes (P-100, P-60, P-40), yet a trend is notable in microcosms containing small protozoa (P-10 and P<10). (b) Boxplots display the alpha diversity parameter at the end of the experiment (96h). Differences between the groups was assessed using an aligned rank transformed ANOVA (ART) procedure and *post hoc* test ART-C test, with different letters above the boxes signifying significant differences between the groups.

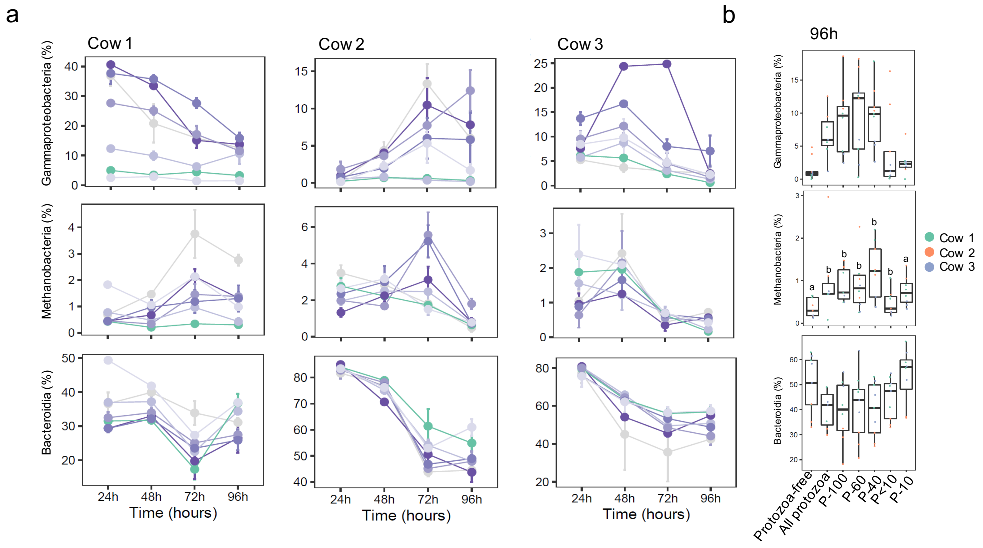


| Factor | P-value |
|---|---|
| Cow | $1.809 \times 10^{-6}$ |
| Protozoa fraction | $1.253 \times 10^{-7}$ |
| Cow x Protozoa fraction | 0.0426 |

| Factor | P-value |
|---|---|
| Cow | 4.71E-06 |
| Protozoa fraction | 1.61E-05 |
| Cow x Protozoa fraction | 0.071 |

| Factor | P-value |
|---|---|
| Cow | $1.387 \times 10^{-6}$ |
| Protozoa fraction | 0.000986 |
| Cow x Protozoa fraction | 0.709 |

**Supplementary Figure S5. Relative abundance of taxonomic classes in the microcosms over time.** (a) Relative abundance of class level taxa. Shown are dynamics of abundance across the different protozoa communities inoculated in the microcosms throughout the experiment (every 24h). (b) Boxplots display the relative abundance at the end of the experiment (96h). Differences between the groups was assessed using aligned rank transformed ANOVA (ART) procedure and *post hoc* test ART-C test, with different letters above the boxes signifying significant differences between the groups.

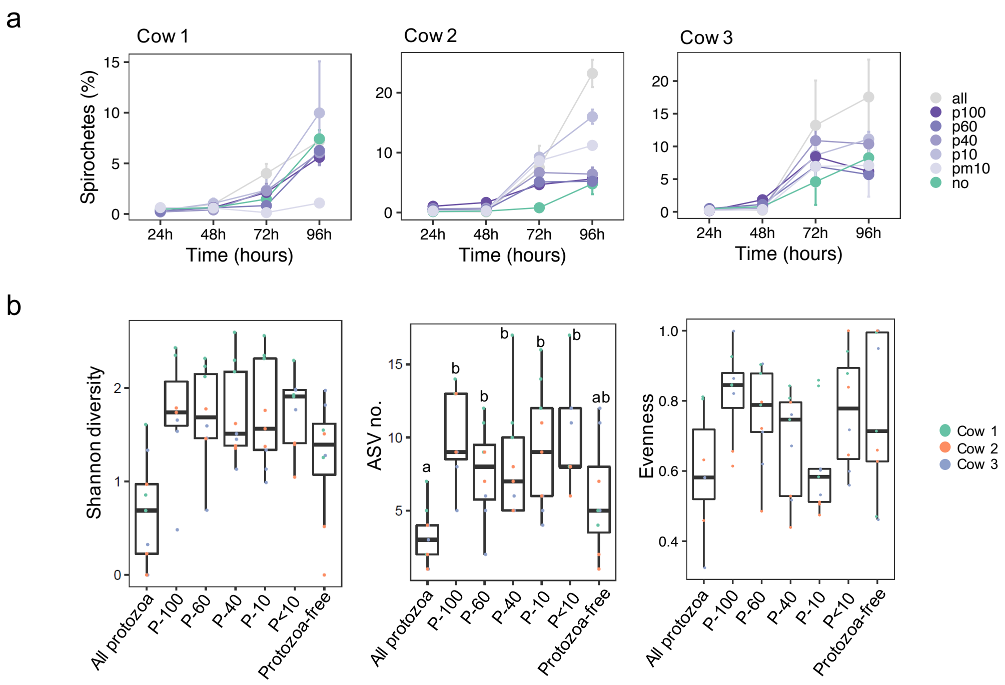


**Supplementary Figure S6. Relative abundance and diversity parameters of the genus *Treponema*.** (a) Relative abundance of the genus Treponema across time. This genus exhibited an increase in abundance across time, which was not significantly different between the protozoa-free microcosms and most protozoa containing microcosms except in the all-protozoa and P<10 microcosm. (b) Shannon diversity, ASV richness (ASV no.) and evenness of the genus *Treponema* at the end of the experiment (96h) is displayed across the different microcosms. We note a significantly higher Shannon diversity and ASV no. in all the groups containing protozoa. Community evenness is a metric that is strongly tied to the number of ASVs, particularly in cases when only one ASV is observed which would lead to an evenness value of 1. This only happened in the protozoa-free communities as these were the only ones observed to have a single Treponema ASV. Differences between the groups was assessed using aligned rank transformed ANOVA *post hoc* test ART-C, with different letters above the boxes signifying significant differences between the groups.

**Supplementary Tables**

**Supplementary Table S1.** Statistical analysis of metabolites shown in Fig.1d and S1.

| | *P*-value | | | |
|---|---|---|---|---|
| **Metabolites** | **24h** | **48h** | **72h** | **96h** |
| Methane | 0.0007 | 0.000033 | 0.0002 | 0.21 |
| VFAs | 0.33 | 0.007 | 0.0441 | 0.0408 |
| Acetate | 0.4 | 0.00013 | 0.001313 | 0.05419 |
| Propionate | 0.56 | 0.11 | 0.3702 | 0.4118 |
| Butyrate | 0.077 | 0.0003 | 0.0755 | 0.09576 |
| Valerate | 0.5 | 0.38 | 0.6524 | 0.5244 |
| Isovalerate | 0.09 | 0.008 | 0.001441 | 0.01223 |
| Isobutyrate | 0.21 | 0.028 | 0.003348 | 0.4429 |

**Supplementary Table S2.** Linear regression of protozoa size (shown Fig.1c) against the produced metabolites over time.

| **Metabolite** | **Time** | $R^2$ | ***P*-value** |
|---|---|---|---|
| Methane | 24 h | 0.003771 | 0.1752 |
| | 48 h | 0.1469 | 0.061848 |
| VFA | 24 h | 0.02376 | 0.168 |
| | 48 h | 0.2631 | 0.0028752 |
| Acetate | 24 h | 0.03051 | 0.1409 |
| | 48 h | 0.3671 | 0.00012896 |
| Propionate | 24 h | 0.008446 | 0.254 |
| | 48 h | 0.1637 | 0.0404 |
| Isobutyrate | 24 h | 0.01726 | 0.1996 |
| | 48 h | 0.2133 | 0.011128 |
| Butyrate | 24 h | 0.06953 | 0.05281 |
| | 48 h | 0.2423 | 0.0051056 |
| Isovalerate | 24 h | 0.1425 | 0.008634 |
| | 48 h | 0.2164 | 0.010264 |
| Valerate | 24 h | -0.01875 | 0.6103 |
| | 48 h | 0.01885 | 0.1913 |
